# Deciduous forests hold conservation value for birds within South Andaman Island, India

**DOI:** 10.1101/2024.04.30.591807

**Authors:** Arpitha Jayanth, Zankhna Patel, Mohammed Mubeen, M Karthikayan, Rohit Naniwadekar

**Author notes:** **Corresponding author:** Arpitha Jayanth. **Open Research statement:** Data and codes used in this study will be uploaded on Zenodo upon acceptance of this manuscript.

## Abstract

Greater diversity of habitats on islands is often correlated with higher species richness (including endemic and threatened taxa), implying the need to understand species-habitat associations. Such habitat associations would also point towards the role of abiotic filtering and competition in structuring species communities, necessitating the examination of the role of species traits and phylogenetic relationships in intra-island community organisation, an aspect poorly examined in literature. We investigated the composition and structuring of forest bird communities in closely co-occurring evergreen and deciduous forests within South Andaman Island (Indian Ocean), wherein the importance of deciduous forests for birds is undervalued. We sampled 27 transects over two years and compared bird species composition and diversity across the two habitats. We examined species-specific associations with habitat (forest) type, basal area, and distance from human settlements, and tested if these responses were explained by species functional traits and tested for phylogenetic niche conservatism after factoring the effects of environmental predictors. Bird species compositions were markedly distinct across the two habitat types, with deciduous forests having greater taxonomic, but not functional or phylogenetic, diversity of forest birds. The distribution of forest birds, including several endemic and threatened species within the island was largely explained by habitat type (with 39% of the bird species analysed showing higher occurrence probabilities in deciduous forests), followed by distance from human settlements and basal area. We did not find evidence of species traits or phylogenetic relationships mediating these habitat preferences, perhaps due to a relatively impoverished species pool, as is typical on islands. Nevertheless, our results underscore the value of deciduous forests in harbouring high islandic species diversity, and being the preferred habitat of several endemic and threatened bird species. Given the historic focus on evergreen forests and the increasing anthropogenic pressure on the forests of the Andamans, we highlight the critical need to include the rapidly diminishing deciduous forests in existing conservation plans.

## INTRODUCTION

Islands have long been a focal interest for developing and testing theories of ecological and evolutionary processes (Whittaker et al. 2017). They are critical ecosystems, being hotspots of unique and endemic biodiversity, as well as hotspots of biodiversity loss (Russell and Kueffer 2019, Matthews et al. 2022). Over 95% of global bird and mammal extinctions have occurred on islands (Loehle and Eschenbach 2012). Island species-area relationships (SARs) generally show that larger islands harbour a greater number of species (Triantis et al. 2012), potentially making them important candidates for the conservation of island biodiversity. Most island biogeography and biodiversity studies focus on patterns and processes between different islands, while processes that maintain diversity within a given island are relatively overlooked (but see Sfenthourakis and Panitsa 2012). In many island systems, SARs have been demonstrated to arise from increased habitat diversity within islands (‘habitat diversity hypothesis’; Kallimanis et al. 2008, Ricklefs and Lovette 1999, Hortal et al. 2009). This emphasises the need to understand the role of different habitat types in maintaining high species richness within islands.

Species compositions can vary across habitats that co-occur within an island, leading to greater intra-island species diversity (Sfenthourakis and Panitsa 2012, Chiarucci et al. 2021). This implies that species can be associated with, or show preference for certain habitat types, even within an island. Birds are highly mobile, able to colonise and perform critical ecosystem functions on oceanic islands. Within islands, bird species assemblages and habitat preferences have been found to differ across habitats that span gradients of vegetation structure, composition, and anthropogenic disturbances (Freifeld 1999, Díaz et al. 2005, Trainor 2005). However, the role of ecological mechanisms like filtering, competition and niche conservatism in structuring intra-island bird communities has seldom been examined.

Bird morphological traits, which are influenced by phylogenetic affinities, are closely linked to their niches (Pigot et al. 2020). Compositional differences between bird communities across habitats could either arise through niche-mediated processes, such as habitat filtering or competition, or from idiosyncratic species responses to habitat characteristics. Clustering of species with similar traits or closer phylogenetic relatedness within habitats could suggest the role of habitat filtering, while overdispersion of traits or phylogeny might suggest the role of competition in mediating bird habitat preferences (Cadotte et al. 2013). These processes have been found to structure bird communities within archipelagos (Si et al. 2017, Triantis et al. 2022, Matthews et al. 2023). However, surprisingly few studies have attempted to understand the role of species traits and phylogenetic relatedness in structuring bird communities, or mediating habitat preferences within islands (but see Johnson 2023, for plants). Although species traits have been found to explain intra-island habitat segregation between native and exotic birds (Barnagaud et al. 2022), and shape bird species co-occurrences at the global scale (Barnagaud et al. 2014, Sato et al. 2020), the role of traits and phylogeny in mediating fine-scale habitat preferences of native and endemic species within islands remains unclear. Moreover, from a conservation point of view, comparing bird communities using functional and phylogenetic metrics could shed light on the importance of certain habitats for functionally or evolutionarily distinct species.

The Andaman Islands (Indian Ocean) lie in the Indo-Burma biodiversity hotspot and are home to endemic and unique tropical flora and fauna. Twenty-one bird species in the Andamans are island endemics, out of which five are categorised as threatened under the IUCN Red List. Larger islands in the Andaman archipelago are deemed critical for bird conservation as they sustain higher species richness of birds, and harbour rarer species that are often absent on smaller islands (Davidar et al. 2001, Gooriah et al. 2020). Contrastingly, protected areas in the Andaman archipelago are greatly skewed towards smaller islands (Davidar et al. 1995), while most forests on larger islands face pressures from selective logging, land diversion, and urbanisation (Sekhsaria 2004, Reddy et al. 2016). Furthermore, large islands have a greater diversity of habitat (forest) types, a key influence on species richness in the archipelago (Davidar et al. 2001). Despite the relevance for bird conservation, the role of different habitat types and disturbance towards the structuring of bird communities, and the distribution of endemic, functionally, and evolutionarily distinct species within larger islands remains unclear.

The forests of the Andaman Islands co-occur as mosaics of evergreen and deciduous patches, with distinct plant species compositions and phenologies (Parkinson 1923, Surendra et al. 2021). Previous examination of habitat preferences of Andaman forest birds has had limited sampling effort or is restricted to specific groups, such as raptors (Thiollay 1997, Yoganand and Davidar 2000). Based on these studies, conservation recommendations have primarily focussed on protecting evergreen forests in the archipelago (Davidar 1996, Davidar et al. 2010), while we know little about the contribution of deciduous habitats towards island bird diversity. Moreover, abiotic conditions and resources available in evergreen and deciduous habitat types likely differ, presenting an interesting opportunity to examine the influence of traits and phylogeny in mediating preferences to such contrasting habitats and species-specific responses to disturbances within an island.

Given the importance of large islands with diverse habitats in maintaining high bird species richness, we explore the intra-island structuring of forest bird communities across two distinct forest habitats, and elucidate the conservation value of evergreen and deciduous habitats for birds in the Andaman Islands. Specifically, at the community-level we (a) compare forest bird composition and taxonomic diversity across evergreen and deciduous habitats, and (b) evaluate standardised mean pairwise distances for functional traits and phylogeny, to detect signals of potential niche-based processes (habitat filtering or competition) in the two habitats. We then (c) analyse species-specific responses to habitat type, habitat structure and disturbance, (d) determine the role of species traits in mediating these responses, and (e) examine the role of phylogenetic niche conservatism in species’ responses. We sampled across South Andaman Island (SA), which is one of the largest islands in the archipelago, home to the administrative capital, and therefore faces rapid urbanisation and development. Apart from the Jarawa Tribal Reserve, the only terrestrial protected area exempt from logging in SA is Manipur Parvat (erstwhile, Mt Harriet) National Park, which consists primarily of evergreen forests. All other evergreen and deciduous forests on the island are subject to logging, extraction, and development. Our study provides species-wise insights on the structuring of native and endemic bird communities, and is informative of the conservation value of different forest types within an understudied and vulnerable tropical island.

## MATERIALS AND METHODS

### Study site

The study was conducted on South Andaman Island (11°07’ – 12°15’ N, 92°30’ – 92°50’ E), located in the Andaman archipelago in the Indian Ocean (Figure 1). The island is one of the largest in the archipelago, covering an area of about 1500 km^2^ and is part of the Indo-Burma biodiversity hotspot. The elevation ranges from 0 – 360 m above sea level on the island, and is unlikely to be relevant in structuring bird communities. The vegetation primarily consists of *Dipterocarpus*-dominated evergreen and semi-evergreen forests, *Pterocarpus*-dominated deciduous forests and littoral forests (Parkinson 1923). The forests of the Andaman Islands have a long history of intensive logging and timber extraction, dating back to the 1800s (Sekhsaria 2004). There is now a legal ban on logging in certain parts of the island (Protected Areas and indigenous Tribal Reserves). Other forested areas (Reserved Forests) continue to be selectively logged, albeit with extraction limits similar to reduced-impact logging practices, specified in the Working Plans of the Andaman and Nicobar Forest Department (A&N Forest Department) (Singh 2003, Mallick 2016, Surendra et al. 2021).

**Figure 1.**
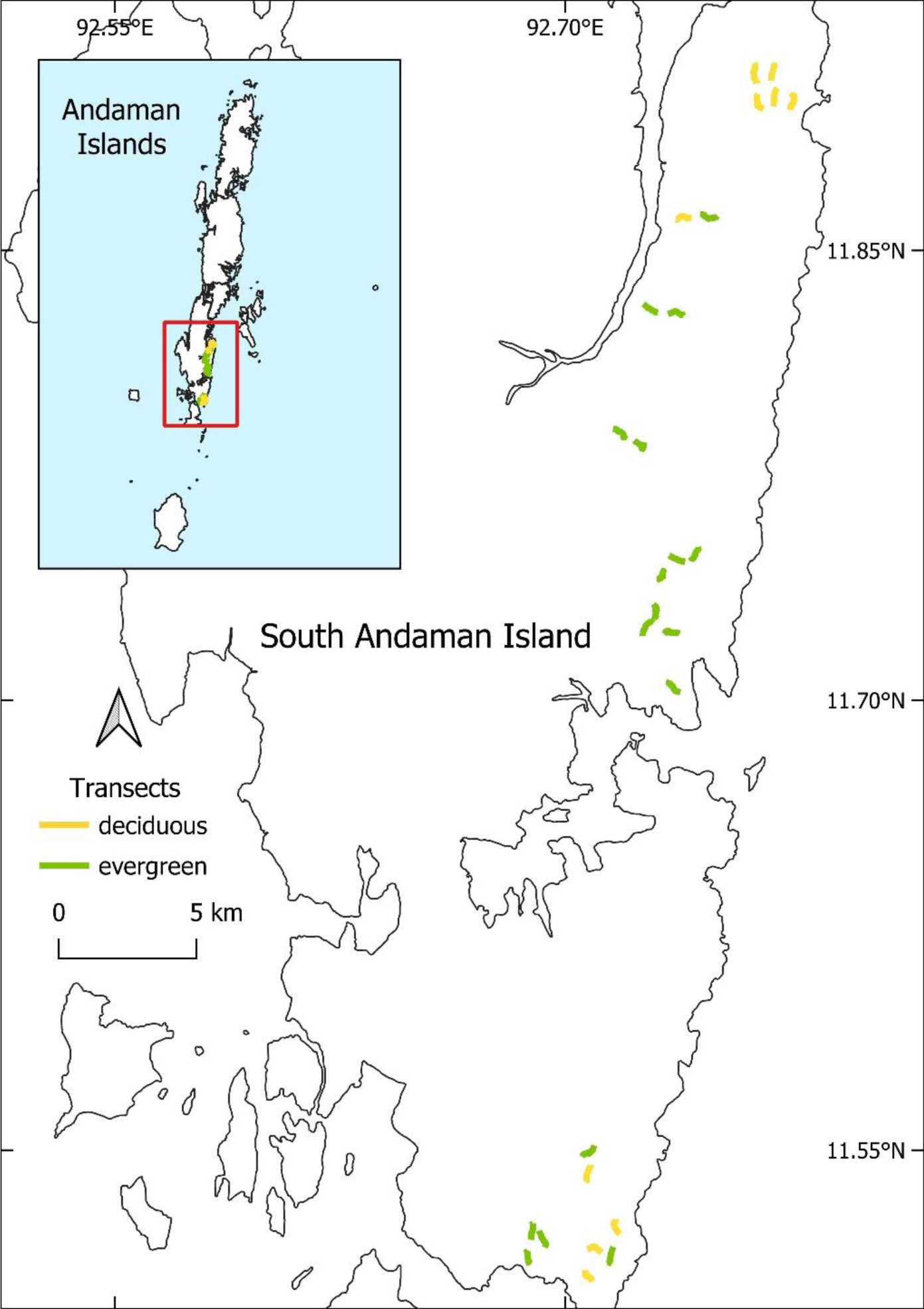
Map of study area in South Andaman Island depicting transects in evergreen and deciduous habitats. The inset shows the location of the study area (red box) within the Andaman archipelago.

We sampled in the forests of Manipur Parvat National Park (erstwhile Mt Harriet National Park) and the Reserved Forests of Shoal Bay and Chidiyatapu. Using forest type classification maps (A&N Forest Department), on-field observations and past logging records by the Forest Department (Singh 2003, Mallick 2016), we identified evergreen and deciduous forest patches, which differed in the time they were last reported to be logged. The forests of the National Park have not been logged since 1979, while some of the selected patches in the Reserved forests have been last logged in the past decade. Within these identified forest patches, we marked 27 transects of length 500 m each (Figure 1). We ensured a minimum distance of 500 m between transects to minimise spatial autocorrelation.

### Sampling bird communities

We conducted transect surveys to record forest bird composition across habitat types and logging histories, from March to May 2022 (Year 1), and from December 2022 to April 2023 (Year 2). In Year 1, all 27 transects were sampled seven times each, and every transect was walked at least two days apart, alternating between morning and afternoon walks. In Year 2, a subset of 10 transects (five evergreen and five deciduous) were sampled eight times each and only in the morning. The total sampling effort across the two years was 134.5 km.

Two observers walked each transect either in the morning (between 0515 – 0930 hrs) or in the afternoon (between 1515 – 1730 hrs). For every bird detected, we noted species identity, group size (if visible), time of detection, and whether the bird(s) was seen or heard. For the analyses, we excluded all detections of flying birds (since they may or may not be using the habitat associated with the transect), and those detected beyond 70 m on either side of the transect (due to higher uncertainty of distance estimation beyond this distance by observers). Any detections of nocturnal species (owls and nightjars) were excluded from the analyses.

### Habitat variables and measuring extraction

To examine responses of different bird species to habitat type and alteration of habitat due to logging and extraction, along each transect we quantified habitat structure, habitat type (proportion of deciduous trees), amount of woody plant extraction, and checked for indicators of disturbance, as described below.

We measured habitat structure using the point-centered quarter (PCQ) method (Cottam and Curtis 1956), during the dry season (March – April 2022) when the deciduous canopy had mostly shed its leaves. Along each transect, we marked 11 equi-spaced points (50 m apart) and estimated tree density and basal area (per hectare) for each transect. We measured canopy cover at each of these points using a canopy densiometer (Spherical Crown Densiometer – Model A). We averaged the canopy cover across the 11 points for each transect. For every tree measured in the PCQ method, we recorded if it was evergreen or deciduous (based on the presence and colour of mature leaves). The proportion of these 44 trees that were deciduous was taken as the proportion of deciduous trees for a given transect. Within a 10 m radius of each PCQ point, we noted the presence of cane, bamboo runners and clumps of standing bamboo as possible indicators of disturbance (Larpkern et al. 2011). We calculated the proportion of points in each transect with the above three types of vegetation.

Forest Department Working Plans are informative on where and when logging had taken place, along with details of timber extracted from the area. However, the habitat may further be modified through damage to non-target individuals during logging, as well as unauthorised extraction of various woody species. To arrive at more precise levels of extraction, we counted the number of all visible cut stumps (similar to Khai et al. 2016) within 20 m-wide belts along each transect. We measured the girth at breast height (GBH) of each stump at the point where it was cut, and later classified these as the number of cut poles (10 cm ≤ GBH < 30 cm) and number of cut adult trees (GBH ≥ 30 cm) per transect. Stumps from relatively older logging events may not be detectable due to higher decomposition rates in tropical moist forests. To detect past logging signatures and to avoid underestimating the same, we counted the number of large trees (GBH ≥ 150 cm) in the 20 m belts. The assumption here is that the number of large trees might be lower in areas that have faced high logging pressures in the past. We also measured the distance (in m) of each transect from the edges of the nearest settlement/village using Google Earth Pro (version 7.3.6.9750) as a proxy for potentially unmeasured human disturbances.

### Statistical analysis

#### Categorisation of transects into habitat types

To reduce subjectivity in classifying transects into evergreen and deciduous habitat types, we used cluster analysis using the partitioning-around-medoids approach (PAM) (Kaufman and Rousseeuw 2009) on transect-level habitat variables. The optimal number of clusters was two, which was determined using the silhouette algorithm. Therefore, we used two a-priori clusters on a Gower-distance measure of the transect-level habitat variables – tree density, basal area, canopy cover, proportion of deciduous trees, number of large trees, number of cut poles, number of cut adult trees, proportion of cane, and proportion of bamboo runners. Categorisation of the transects into these two clusters closely matched our initial categorisation into evergreen and deciduous habitat types on field. The clustering was validated by computing the silhouette width i.e., average distance between clusters (values close to 1 indicate that the object is well clustered). Due to the large number of habitat variables involved, we initially ran a principal component analysis (PCA) on the above. Only 57% of the variation was explained by the first two axes (Figure 2a), while 93% of the variation was explained by six axes. We used R packages ‘cluster’ for the medoid clustering, ‘FactoMineR’ and ‘factoextra’ for the PCA, and ‘tidyverse’ for data processing.

**Figure 2.**
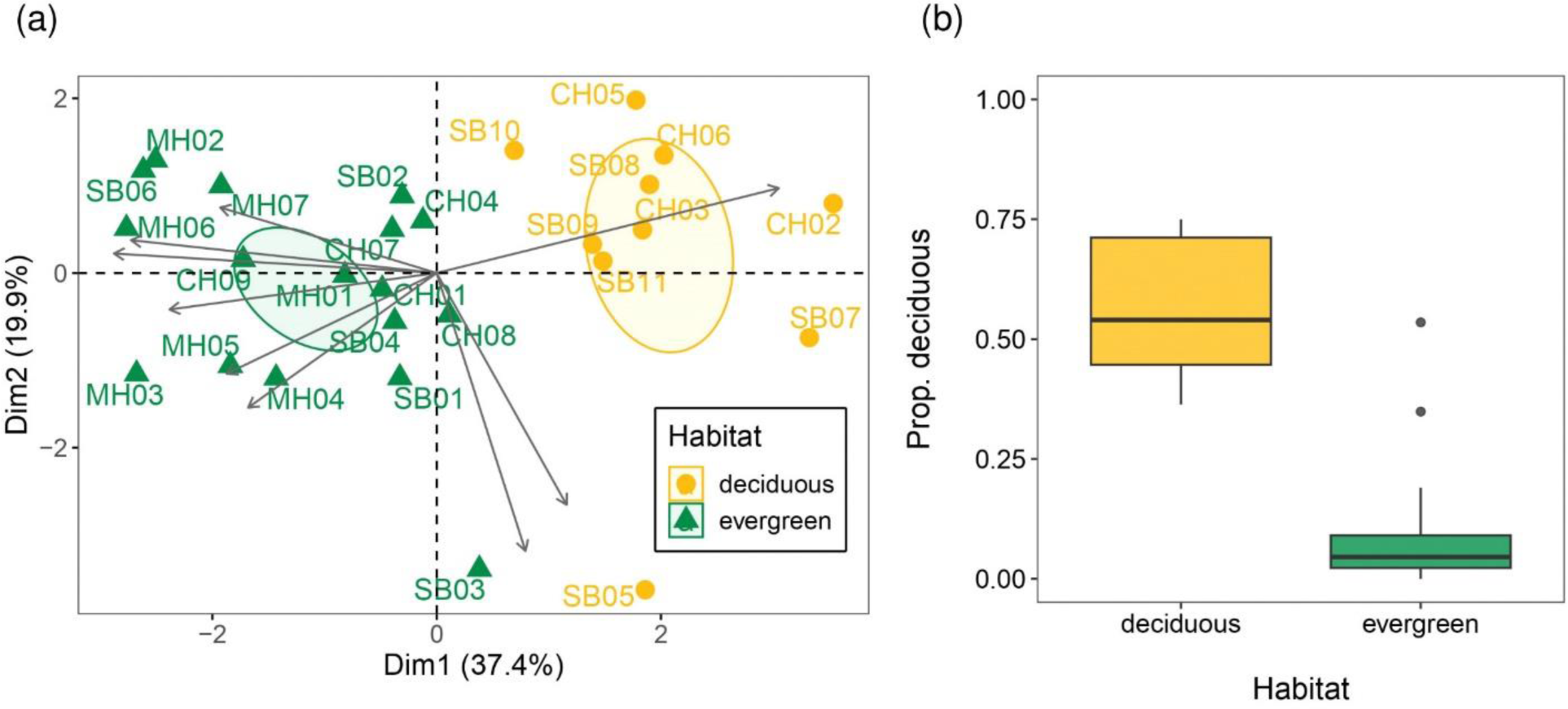
(a) Biplot from the principal component analysis (PCA) depicting the first two axes, along with habitat and extraction-related variables. The first two axes of the PCA explain only about 57% of the variation in habitat and extraction levels between transects. Points represent the 27 transects across the study site. Grey arrows represent different habitat variables – clockwise from top-right quadrant: proportion of deciduous trees, number of cut poles, number of cut adults, proportion of points with bamboo runners, proportion of points with cane, basal area (per ha), average canopy, tree density (per ha), number of large trees. The two habitats have been depicted using 95% confidence ellipses, based on the PAM clustering results. (b) Boxplot depicting proportion of deciduous trees (sampled using the PCQ method) in evergreen and deciduous habitats.

#### Bird community composition across habitat types

To test for differences in bird species composition across evergreen and deciduous habitat types, we used average detections of each species per transect (to account for the imbalanced effort per transect) and constructed a distance matrix using the Bray-Curtis dissimilarity index. We visualised this on a Non-Metric Multidimensional Scaling (NMDS) plot using the first two axes. We computed the difference in bird community composition across evergreen and deciduous clusters using the R function *anosim*, where a value close to 0 indicates that the two communities are similar and a value close to 1 indicates that the two communities are different. We used the R package ‘vegan’ for this analysis.

#### Bird diversity across habitat types

Since we had unequal sampling effort in evergreen (79.5 km) and deciduous (55 km) habitats, we used sampling coverage to evaluate our sampling completeness. Sampling coverage estimates the proportion of individuals in the community that belong to the set of species already detected in the sample (Roswell et al. 2021). Our sampling coverage was >99% for both habitat types. We pooled species detections across temporal replicates and standardised coverage for both habitats at 99%. Using the R package ‘iNEXT’, we computed Hill-Shannon diversity and compared across the two habitats. Hill-Shannon measures are ideal for diversity comparisons since rare species are given less emphasis and standardisations are found to be robust (Roswell et al. 2021). We bootstrapped the data 50 times to arrive at 95% confidence intervals. We interpreted statistically detectable differences in species diversity if the confidence intervals did not overlap, following (Cumming et al. 2007).

#### Functional and phylogenetic structuring across habitat types

We used mean pairwise distances (MPD) to determine if bird communities across habitat types were functionally and phylogenetically clustered or overdispersed, following (Chapman et al. 2018). We used average detections of each species per transect to account for the unequal sampling effort. For each transect, we computed functional MPD (fMPD) using a distance matrix of species traits related to morphology (Tobias et al. 2022) and diet (Wilman et al. 2014); and phylogenetic MPD (pMPD) using a time-calibrated phylogenetic tree from birdtree.org (Jetz et al. 2012). Details of functional trait selection are given in Appendix S1: Figure S2. We compared standardised effect sizes (ses) of fMPD and pMPD to evaluate clustering or overdispersion across the two habitats. We calculated sesMPD values using the function *ses.mpd* in the R package ‘picante’, with an independent swap algorithm on the island-level species pool, which generated null communities by weighting the random drawing of a species by its abundance. By creating null communities from the island-level species pool instead of mainland or archipelago-level pools, we were able to exclude any effects of filters influencing establishment or dispersal to the study island.

#### Species-specific associations with habitat type and disturbance

We examined associations of bird species with habitat structure, habitat type and disturbance using the Hierarchical Modelling of Species Communities (HMSC) approach (Ovaskainen and Abrego 2020). This is a class of joint species distribution model that tests species responses to habitat or environmental variables, examines if these responses can be explained by species traits, and checks for any phylogenetic signal in the residuals. Here, we used each transect walked as a sampling unit (N = 269). For 33 bird species (detected on >5% of transect walks), we modelled occurrence probability using a presence-absence model with a probit link. Species responses were modelled as a function of basal area (habitat structure), proportion of deciduous trees (habitat type), and distance to the nearest settlement (proxy for disturbance). We analysed if these species responses were mediated by their traits. For this, we used species traits related to diet (proportion of fruit, seed, and nectar in the diet, obtained from Wilman et al. 2014) and morphological measurements related to foraging (beak width) and locomotion (tarsus length, tail length and Kipp’s distance) taken from Tobias et al. (2022), in the analyses. Please see Appendix S1: Figure S3 for details on the selection of habitat variables and traits. We used a time-calibrated phylogenetic tree from birdtree.org (Jetz et al. 2012) to determine any phylogenetic signal in the residual variation of species responses to the habitat variables mentioned above. Spatial coordinates of the transects and temporal sampling replicate were used as random effects, to test for spatial and temporal autocorrelation. The HMSC model was fitted with the R package ‘Hmsc’. We sampled the posterior distribution with three Markov Chain Monte Carlo (MCMC) chains, and removed the first 50,000 iterations as burn-in. The chains were thinned by 1000 to arrive at 250 samples per chain. We assumed MCMC convergence if the potential scale reduction factors of the model parameters were <1.05. We evaluated model fit for the presence-absence model using Tjur’s *R^2^*. We interpreted a habitat (or trait) variable to have a statistically detectable effect on bird species occurrence only if the 95% credible intervals for the estimated β (or γ) coefficients for the predictor did not overlap zero.

## RESULTS

### Habitat classification

The transects were categorised into two a-priori clusters, or habitat types (17 as evergreen and 10 as deciduous; Figure 2a). The average silhouette distance between the two clusters was 0.33, indicating that the transects were moderately well categorised (Figure S1). The two clusters/habitat types differed in the proportion of deciduous trees (Figure 2b).

### Bird community composition and structuring across habitat types

We recorded a total of 9,921 detections of birds (both visual and auditory) from 54 species. Species composition of birds in the two habitat types was statistically detectable and moderately dissimilar (*R*_anosim_ = 0.44, *p* = 0.001, stress = 0.19, Figure 3a).

**Figure 3.**
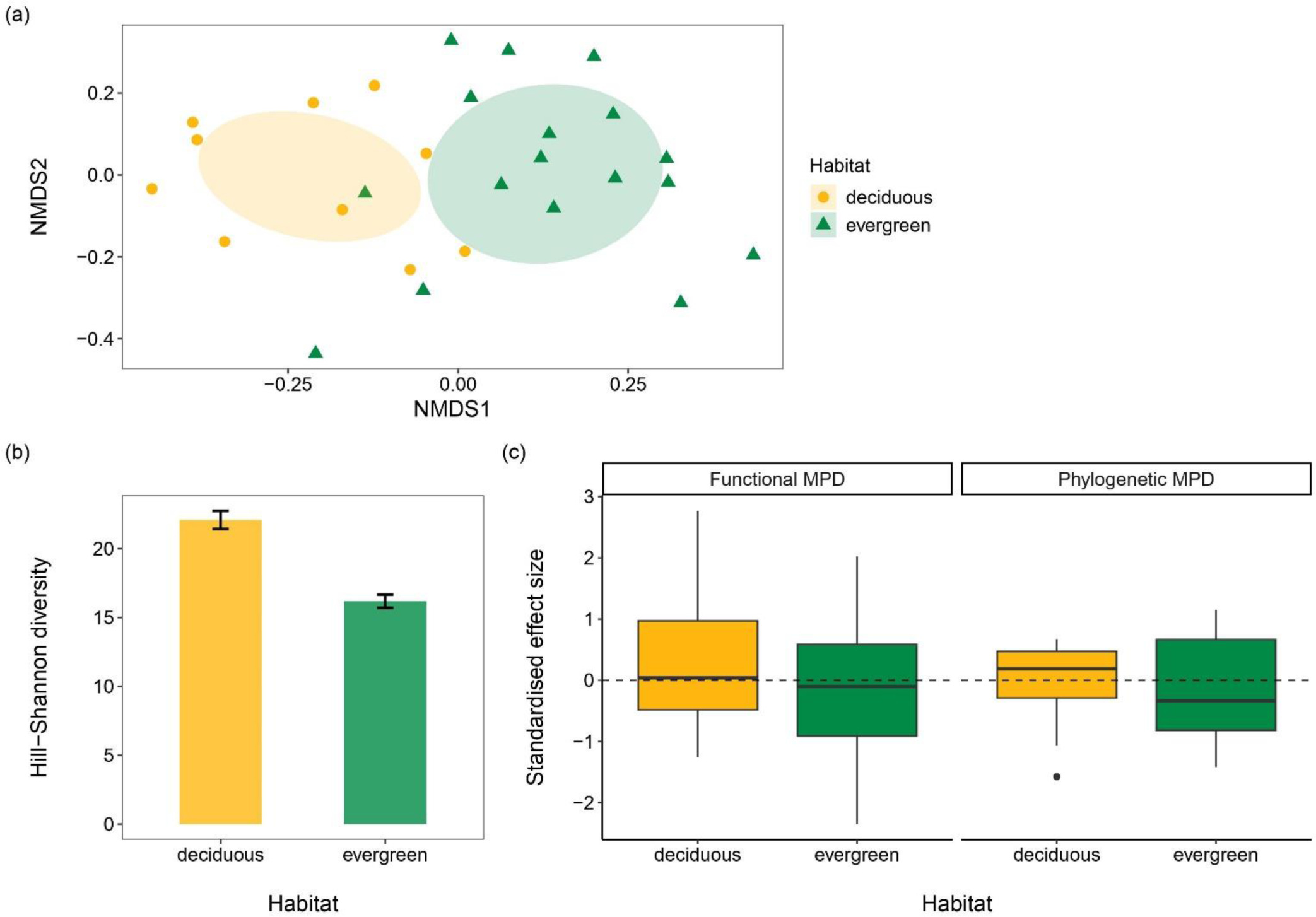
(a) Bird community composition across evergreen (green; filled triangles) and deciduous (yellow; filled circles) habitat types in South Andaman Island. Each transect has been represented as a point, and covariance ellipses have been drawn for each habitat type. An analysis of similarities test (ANOSIM) indicates that bird communities are moderately different across the two habitat types (*R_anosim_* = 0.44, *p* = 0.001). (b) Taxonomic diversity across habitats, compared using the Hill-Shannon measure. Bars indicated bootstrapped 95% confidence intervals. (c) Standardised effect sizes of functional and phylogenetic MPD for each habitat type. Values centred around 0 indicate the lack of clustering or overdispersion in the communities.

Deciduous habitats supported higher taxonomic diversity of bird species than evergreen habitats (Figure 3b). We found no detectable signal of functional or phylogenetic clustering or overdispersion of bird communities in evergreen and deciduous habitat types (Figure 3c).

### Species-specific associations with habitat type and disturbance

The HMSC model sufficiently converged and explained, on average, 11% of the variation across species occurrences for the 33 species analysed. Of the total explained variation, 54.9% was explained by the three fixed effects. Amongst the fixed effects, proportion of deciduous trees accounted for the most amount of explained variation (22.9%) in species occurrences, followed by distance from settlements (18.1%) and basal area (13.9%). Spatial and temporal random effects explained 22.3% and 22.8% of the variation in species responses respectively (Figure 4).

**Figure 4.**
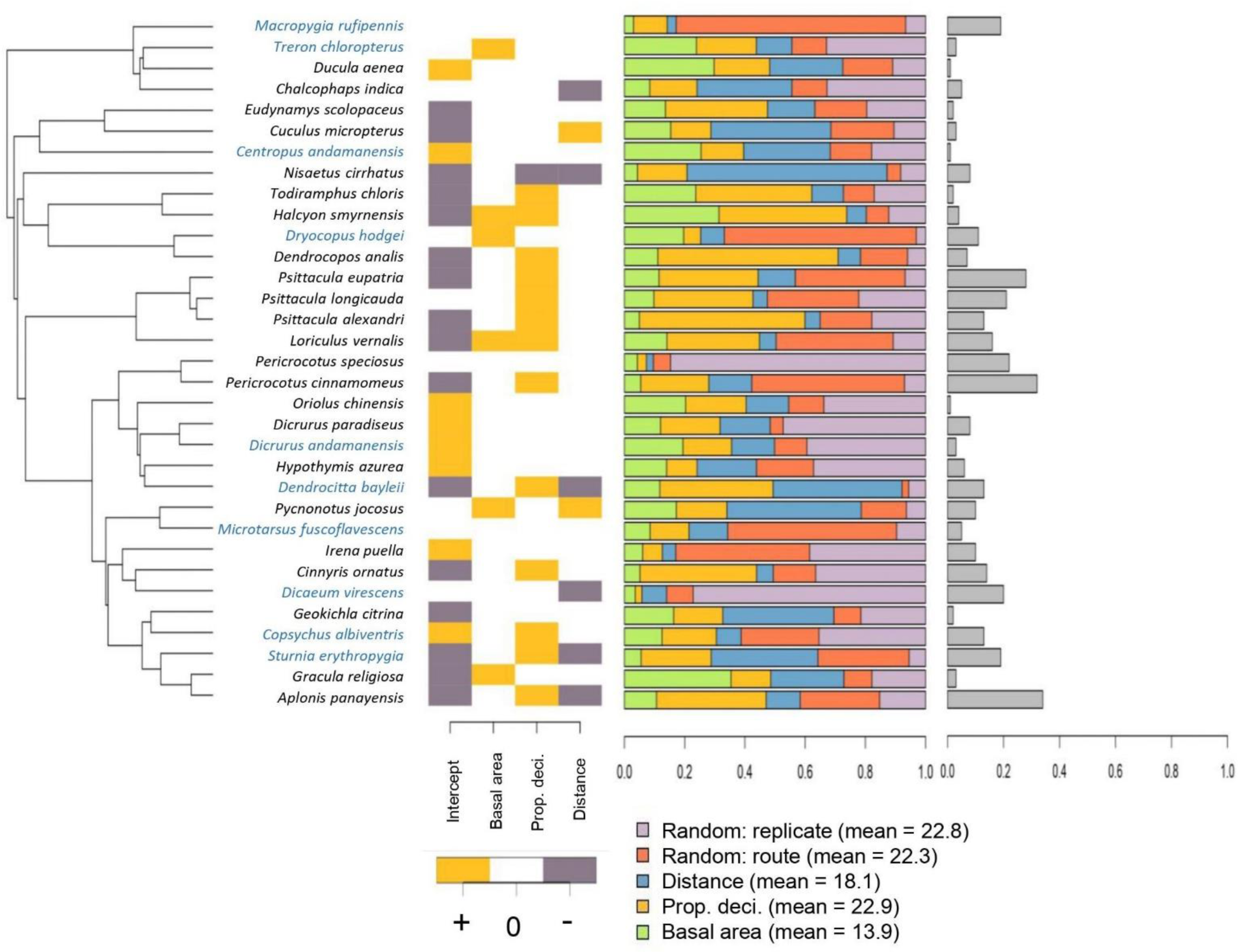
Species-specific responses of birds to habitat variables in terms of their occurrence probability. From left to right: the phylogeny is shown on the extreme left, with tips representing the 33 species of birds analysed (names in blue text indicate endemic species). The response matrix indicates positive responses to habitat variables in yellow and negative responses in purple at ≥ 95% posterior probability. ‘Prop. deci’ refers to proportion of deciduous trees, ‘Distance’ refers to distance from nearest settlement. The variance partitioning plot displays the proportion of *explained* variance that is explained by each of the habitat variables (refer to legend). The bar plots on the extreme right depict the proportion of explained variance (Tjur *R^2^*) in occurrence probability for each species.

The occurrence probabilities for 13 of the 33 bird species analysed (39%) were detectably higher in deciduous habitats, while one species (Changeable hawk-eagle *Nisaetus cirrhatus*) had a lower occurrence probability in deciduous habitats (Figure 4). Of the 13 species associated with deciduous forests, three are endemic (Andaman treepie *Dendrocitta bayleii*, White-headed starling *Sturnia erythropygia* and Andaman shama *Copsychus albiventris*), and four are categorised as threatened by IUCN (Andaman treepie *Dendrocitta bayleii* and Long-tailed parakeet *Psittacula longicauda*: Vulnerable; Alexandrine parakeet *Psittacula eupatria* and Red-breasted parakeet *Psittacula alexandri*: Near Threatened) (Appendix S1: Figure S6). The remaining 19 species showed no statistically detectable preference for evergreen or deciduous habitats (Figure 4).

Six species had higher occurrence probabilities in habitats with higher basal area (Figure 4). Of these, Andaman woodpecker *Dryocopus hodgei* and Andaman green-pigeon *Treron chloropterus* are endemic and classified as Vulnerable by IUCN (Appendix S1: Figure S5).

Geographic distance from settlements positively affected the occurrence probabilities of two species (Indian cuckoo *Cuculus micropterus* and Red-whiskered bulbul *Pycnonotus jocosus*). Six species had higher occurrence probabilities closer to settlements (Figure 4), including endemics such as Andaman treepie *Dendrocitta bayleii* (IUCN Vulnerable), White-headed starling *Sturnia erythropygia* and Andaman flowerpecker *Dicaeum virescens* (Appendix S1: Figure S7).

Species responses to the predictors were not explained by species traits. We did not detect any phylogenetic signal in the residuals (*ρ* = 0.67, 95% CI: 0 – 0.98). We detected some spatial signatures in the model at the scale of approximately 19 km.

## DISCUSSION

Our field study showed that evergreen and deciduous habitats within an island can be quite distinct. We find evidence that these two habitats harbour unique communities of forest birds, with deciduous habitats being more taxonomically diverse. We find that 39% of forest birds (13 of 33 species) show a statistically higher occurrence probability in deciduous habitats, while the occurrences of a few other species were associated with proximity to settlements and/or higher basal area. There was no evidence of functional traits or phylogeny mediating species’ responses or community-level structuring, implying species’ idiosyncratic responses to habitat characteristics. Our results suggest that the currently unprotected deciduous habitats play an important role in maintaining species diversity on the island and are critical for forest bird conservation in South Andaman Island. It is critical that deciduous habitats are included in the Protected Area network on the main island archipelago.

### Conservation value of deciduous forests on larger islands in the Andamans

Diverse habitats within an island are known to harbour distinct species communities (Rodriguez-Estrella et al. 1996, Díaz et al. 2005, Trainor 2005, Katsimanis et al. 2006), thereby contributing to increased diversity at the scale of individual islands (Hortal et al. 2009, Sfenthourakis and Panitsa 2012, Chiarucci et al. 2021). Our analysis of forest bird communities across habitat types *within* a single island reveals clear differences in the composition of bird species between co-occurring evergreen and deciduous habitats. This finding lends explanatory support to the fact that islands with greater habitat diversity in the Andaman archipelago harboured higher species richness (Devy et al. 1998, Davidar et al. 2001) by harbouring species that specialise in respective habitats. This re-iterates the importance of different habitat types for bird diversity within an island. Additionally, habitat type was an important predictor of bird species occurrences in South Andaman Island (Figure 4). When species seem to specialise or prefer certain habitat types within an island, as observed here (Figure 4; and see Yoganand and Davidar 2000), the area available for the persistence of these ‘habitat specialists’ is further reduced. This augments the conservation relevance of diverse habitats with islands and, in the context of the Andamans, necessitates the protection of both evergreen and deciduous habitats.

Previous work from the Andamans Islands highlight the importance of evergreen forests on larger islands for various taxa, including birds (Davidar 1996, Davidar et al. 2001). Our results underscore the value of deciduous forests for at least 39% of forest bird species found here, including endemic and threatened species such as Andaman treepie *Dendrocitta bayleii* and White-headed starling *Sturnia erythropygia* (Figure 4). We also find that deciduous habitats support higher species diversity compared to evergreen habitats (Figure 3b), and are equally important for functionally and evolutionarily distinct bird species on the island (Figure 3c).

However, deciduous forests have received considerably less conservation focus, especially on larger islands like South Andaman. The geographical area of deciduous forests in the Andamans is relatively small compared to evergreen forests (Reddy et al. 2016). While both evergreen and deciduous forests are selectively logged in the islands, some tracts of evergreen forests are given high legal protection and are exempt from felling and local extraction (e.g., Manipur Parvat National Park (Singh 2003, Mallick 2016)). On the contrary, barring patches in the Jarawa Tribal Reserve, most deciduous forests on larger islands are routinely logged and lack the same level of protection, despite being more susceptible to the negative impacts of repeat logging (Khai et al. 2016, Surendra et al. 2021). Over a period of nearly 40 years, loss of forest cover in the Andaman Islands has been estimated to be as high as 21% for deciduous and about 0.2% for evergreen forests, as a result of anthropogenic as well as natural disturbances (Reddy et al. 2016). To conserve bird diversity in the Andaman Islands, we therefore recommend prioritising the protection of deciduous forests, alongside evergreen forests, within large islands like South Andaman.

### Structuring of bird communities within an island

Island species assemblages are the result of a series of historical and ecological processes, often mediated by species traits and phylogeny (Sato et al. 2020). The relative importance of different processes seems to vary with the spatial scale under purview. Most studies examine these processes at the level of archipelagos and islands, and find that species communities at this scale largely exhibit functional and phylogenetic clustering (Si et al. 2017, Baiser et al. 2018, Triantis et al. 2022). This is likely because only a few clades or functional groups can disperse to and colonise isolated islands. At finer scales, for example, within islands or habitats, filtering by environment or habitat could lead to clustering, whereas competition between functionally or phylogenetically similar species could result in overdispersion (Cavender-Bares et al. 2009).

However, empirical evidence for intra-island community assembly is relatively scarce. A study on plants found evidence for clustering due to habitat filtering within islands (Johnson 2023), while dispersal limitation and stochasticity likely led to overdispersion of intra-island microbial communities (Wang et al. 2020). On the contrary, our study detects no signals of community-wide functional and phylogenetic clustering or overdispersion of forest birds between contrasting habitats on the same island (Figure 3c). Although we found that some species and groups (eg: parakeets and sturnids) were associated with one or more habitat variables (Figure 4), our HMSC analysis reveals that neither species traits nor phylogeny explained these responses, similar to Barnagaud et al. (2023). Habitat partitioning between native and exotic birds within islands elsewhere have been found to be trait-dependent (Barnagaud et al. 2022). However, it must be noted that the aforementioned studies compare native and exotic bird assemblages, while our study focusses on habitat differences within the native bird species community. Nevertheless, these discrepant findings suggest that the processes mediating community assembly within islands are perhaps site, taxon and scale dependent, requiring further research to evaluate its pervasiveness and strength. In the case of Andaman forest birds, dispersal to the island possibly imposes a strong filter, likely resulting in diminished functional and phylogenetic diversity within the island in the first place. Within this filtered species pool, any species-habitat associations detected seem to have formed independent of traits and phylogeny.

### Species’ responses to intra-island habitat variables are idiosyncratic

Despite the relatively small area covered by deciduous habitats in our study site, we demonstrate its preference by at least 13 species of forest birds, including several endemic and threatened species. Species groups – such as parakeets and starlings – typically seen in deciduous, secondary growth forests or relatively open areas seemed to have higher occurrence probabilities in deciduous habitats (Figure 4). Asian glossy starlings *Aplonis panayensis* and White-headed starlings *Sturnia erythropygia* were also frequently observed feeding on the fruits of many fleshy-fruited trees in the deciduous habitats (personal observation/unpublished data) and likely play key functional roles as seed dispersers in deciduous and open/degraded habitats. Although we detected positive associations of two species of kingfishers (White-throated kingfisher *Halcyon smyrnensis* and Collared kingfisher *Todiramphus chloris*) with deciduous habitats, our data suggests that their occurrence probabilities are generally low in forested areas (Appendix S1: Figure S6).

Certain species responded to indicators of habitat structure, such as basal area. Higher basal area is generally characteristic of least-disturbed habitats with large, old-growth trees and minimal logging (Khai et al. 2016). Columbid species in the American Samoa have been found to prefer native undisturbed forests, presumably for nesting sites and food resources (Freifeld 1999). In South Andaman Island, two endemics – Andaman woodpecker *Dryocopus hodgei* (Vulnerable) and Andaman green-pigeon *Treron chloropterus* (Near Threatened) – were discerned to have reduced occurrence probabilities in habitats with low basal area. While this was also detected in the case of four non-endemic species, their effect sizes were comparatively small (Appendix S1: Figure S5). This suggests that for some species, especially certain endemic and threatened species, the persistence of habitats with large trees could be important to sustain their populations.

Proximity to human settlements can represent an array of habitats disturbances and alterations, such as canopy openness, edge effects, or higher risk of hunting. In our study site, species detected more often near settlements largely include those that prefer open and/or edge habitats, such as starlings and Andaman flowerpecker *Dicaeum virescens*. In the case of Red-whiskered bulbul *Pycnonotus jocosus*, we statistically found their occurrence to increase away from settlements, contrary to typical patterns observed elsewhere (Goyal et al. 2019). However, a closer inspection reveals that they seem to be generally widespread across gradients of habitats within the island, and their apparent increase with both distance from settlements and basal area seems to be marginal (Appendix S1: Figure S7).

We detected the endemic Andaman woodpigeon *Columba palumboides* (classified as Near Threatened by IUCN) only seven times throughout our sampling. A nationwide assessment of India’s birds also lists the woodpigeon as data deficient, amongst other Andaman endemics (SoIB 2023). Therefore, we recommend a focussed study on this endemic species to determine the size and potential threats to its population in the Andaman Islands.

After accounting for fixed effects, we found that spatial and temporal random effects together explained 41.5% of species occurrences (Figure 4). This implies that species occurrences in South Andaman Island could be additionally influenced by spatially-dependent processes or habitat variables not measured in our study. Certain species distributions may be inherently patchy, owing to territoriality for instance, leading to spatial contingency in their detections. While we have attempted to include major factors influencing bird species distribution (habitat structure, habitat type and proxies for disturbance), further research may investigate the role of territoriality and other biotic interactions in determining species occurrences on the island.

### Conclusions

Our findings emphasise the importance of protecting distinct habitat types within an island to maintain bird diversity in the Andaman archipelago. Both evergreen and deciduous forest types harbour unique communities of birds, and warrant protection from habitat diversion and other disturbances. Our results particularly underscore the value of deciduous forests, which have so far received less conservation attention and experienced significantly higher modification in the Andaman Islands. Apart from habitat type, we also demonstrate that habitat structure and proximity to settlements can influence bird species distribution in the islands, largely independent of niche-based processes. We also highlight the need to examine mechanisms that maintain and structure species diversity within islands. The patchy mosaics of evergreen and deciduous forests in the Andaman Islands form a fascinating system, that together support unique communities of islandic flora and fauna. While we have begun to understand their species compositions and potential alterations to these habitats (this study; Surendra et al. 2021), we are yet to discern the origin and mechanisms through which these distinct habitat types co-occur at fine scales in the Andaman archipelago.

## ACKNOWLEDGEMENTS

We wish to acknowledge Michael (Vela Kujur), Sujith Bengra and Aditya Gadkari, whose contributions towards the execution of this study have been invaluable. We are grateful to the Andaman and Nicobar Forest Department for issuing us the necessary permits (No. F.10(G-I)/39/Vol. XI/985 and No. CWLW/WL/17/Vol-II/1054). We thank Sanjay Waradkar (CF), Nabanita Ganguly (DCF), A.K.Paul (DFO), Soundara Pandian (ACF), Sukumar (ACF), Tilak (ACF), Jagdish (RFO), Mohammed (RFO), Subhashis Ray (RFO), Majeed, Ganeshan, and all the staff of Manipur Parvat National Park and South Andaman Forest Division for their generous support and assistance during our fieldwork. We are grateful to Navendu Page, Sartaj Ghuman, Shashank Dalvi, Evan Nazareth, Elrika D’Souza, Soham Dixit, Sagar Kelkar, Salman Khan, Arun Singh, Nayantara Biswas, Jithin Vijayan, Pooja Pawar, and Andaman Avians Club for support and discussions. We thank Jahnavi Joshi for help with extracting phylogenetic trees and Otso Ovaskainen for clarifying some queries with the HMSC R package. We are especially grateful to the late Josna Mallick, and to Hassan Ali, Dhanalaxmi Kujur, Shiraz Noble, and their families for the generous and warm hospitality during our work in the island.

## Author contributions

AJ and RN jointly conceived, designed, and acquired funding for the study. AJ, ZP, MM and KM collected the data on field. AJ and RN analysed the data and wrote the manuscript. All authors reviewed and approved the final version of the manuscript.

## APPENDIX S1 – SUPPLEMENTARY MATERIAL

**Figure S1.**
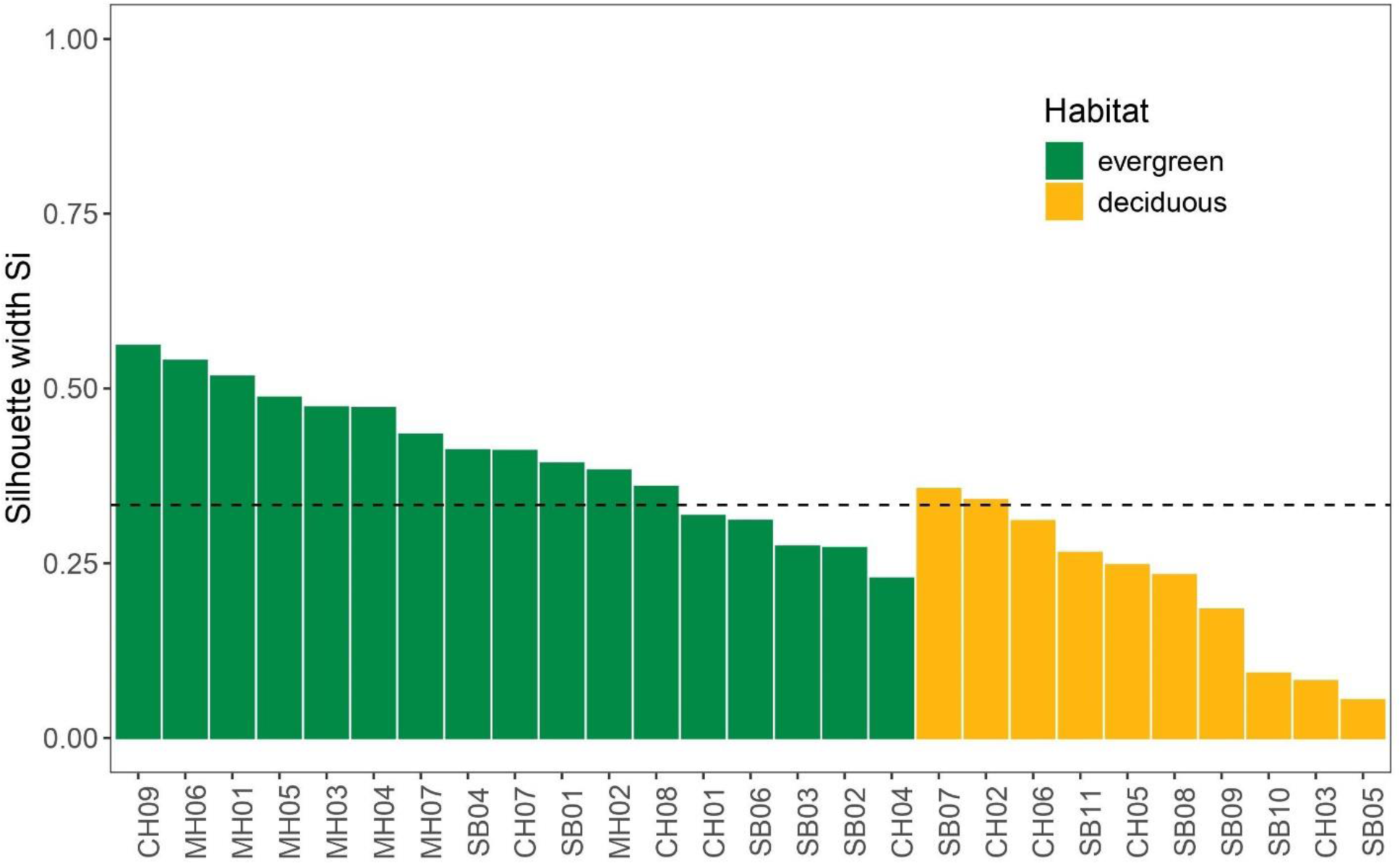
Cluster silhouette plot showing the silhouette width of each transect. Values close to +1 indicate that the transect has been clustered well, while values close to -1 indicate that the transect is probably incorrectly categorised into a given cluster. Green and yellow represent evergreen and deciduous habitat types respectively. The dashed horizontal line indicates the average silhouette width of +0.33.

**Figure S2.**
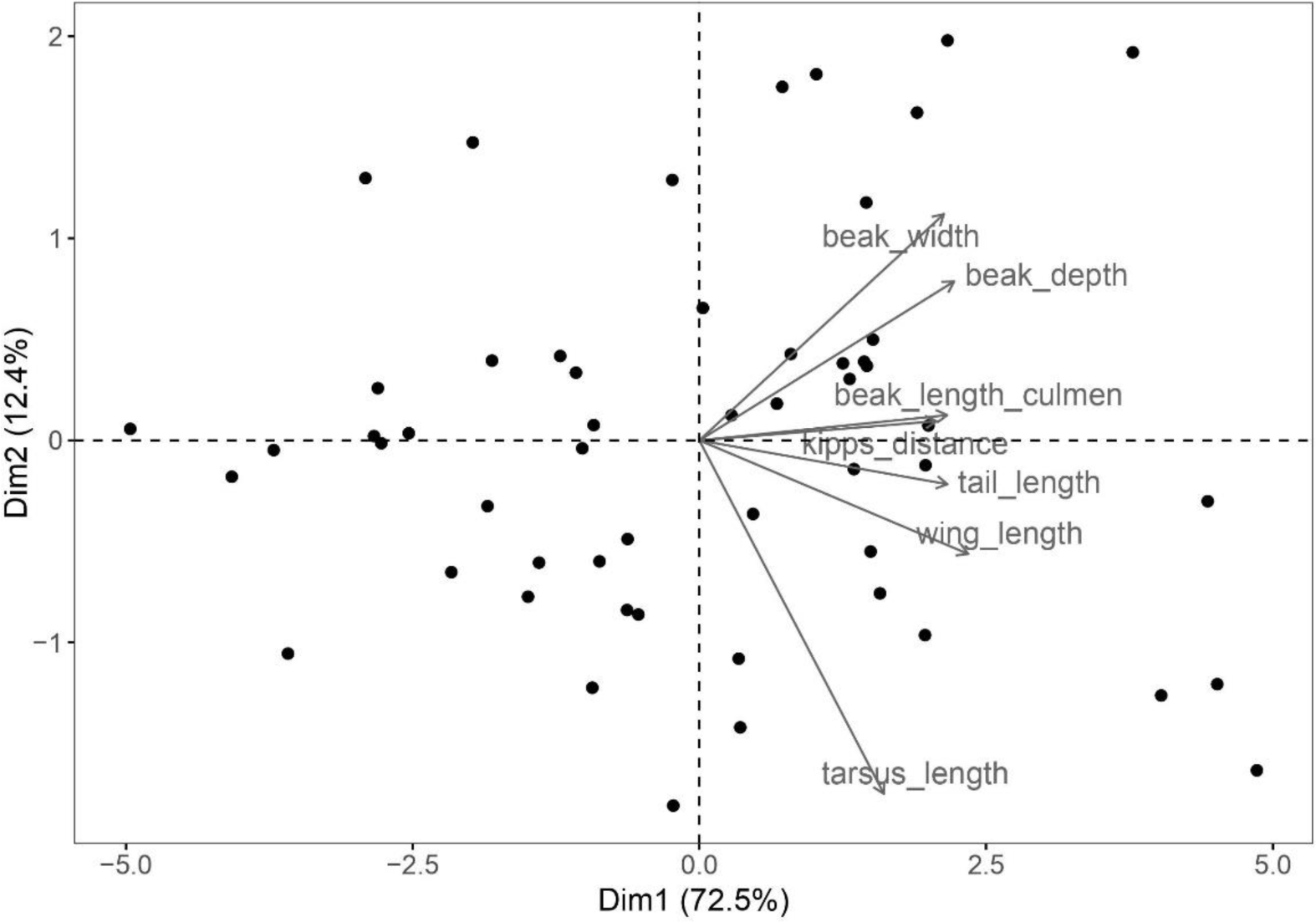
Biplot from the principal component analysis (PCA) done for morphological traits used in the functional diversity analysis. Following (Chapman et al. 2018), we used traits pertaining to trophic niche (beak length, width and depth) and locomotion (tarsus length, wing length, Kipp’s distance, and tail length). The first three axes of the PCA explained 92% of the morphological traits, and were therefore used in the functional diversity analysis. <u>Selection of dietary traits:</u> Similar to morphological traits, we performed a PCA on the proportions of invertebrates, vertebrates, fruit, nectar, seeds, and other plant matter in the diet of each bird species. Five axes of the PCA were required to explain at least 90% of the variation in diet. Therefore, we directly used the dietary proportions listed above in the functional diversity analysis. Since there were relatively few species with seeds and other plant matter in their diets, we combined these into a single dietary variable before proceeding with the analysis.

**Figure S3.**
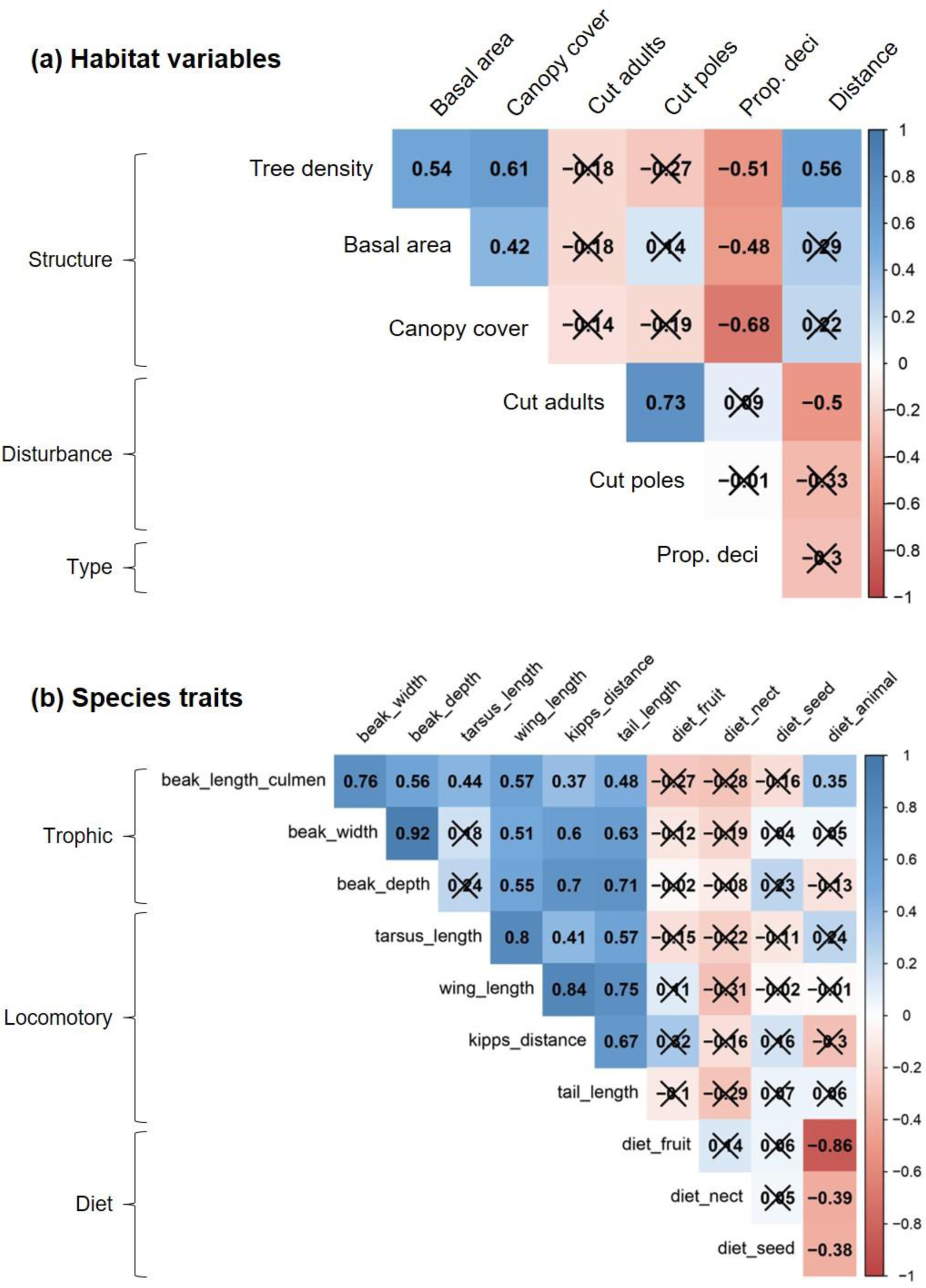
Correlation plots amongst (a) habitat variables and (b) species traits used in the HMSC analysis. The numbers indicate Pearson’s correlation coefficients, and crosses denote correlation coefficients that were statistically not significant (*p* > 0.05). The scale on the right represents correlation coefficient values from negative (red) to positive (blue). <u>Selection of habitat variables (a):</u> We checked correlations between variables capturing habitat structure (tree density, basal area, canopy cover), habitat type (proportion of deciduous trees), and habitat disturbance (number of cut adult trees, number of cut poles, and distance from settlements). To avoid variance inflation due to correlation amongst predictor variables, we used basal area as an indicator of habitat structure, proportion of deciduous trees as habitat type, and distance from settlements as a proxy for disturbance in the HMSC model. <u>Selection of species traits (b):</u> We used traits related to trophic niche (beak length, beak width, and beak depth), locomotion (tarsus length, wing length, Kipp’s distance, and tail length), and diet (proportions of fruit, nectar, seed, and animal matter in the diet), following (Chapman et al. 2018). After excluding variables with high correlation coefficients, we used beak width (trophic niche), tarsus length, Kipp’s distance and tail length (locomotory traits) and proportions of fruit, seed and nectar in the diet as species traits in the HMSC model.

**Figure S4.**
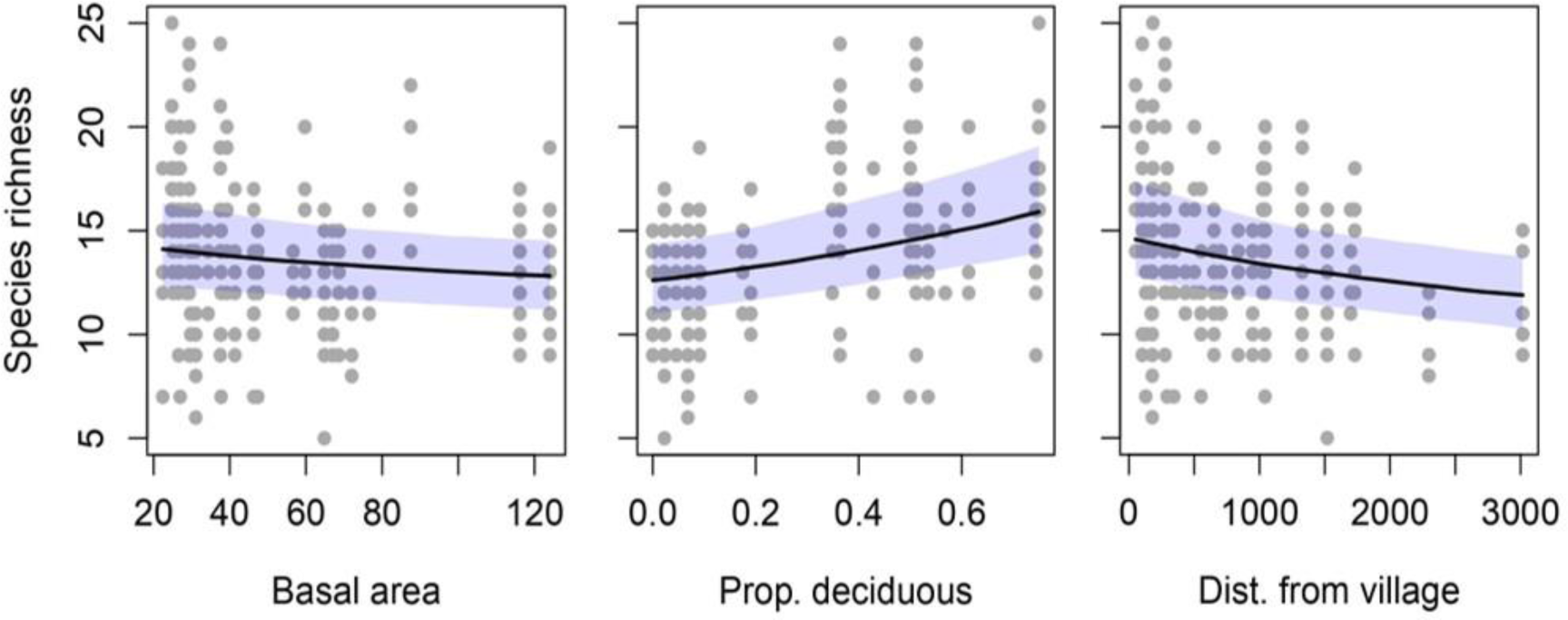
HMSC model predictions of variation in bird species richness along gradients of basal area (per hectare), proportion of deciduous trees, and distance from nearest settlement/village (in m). Black lines represent the posterior median, points represent individual samples, and the shaded area depicts the posterior 95% credible interval.

**Figure S5.**
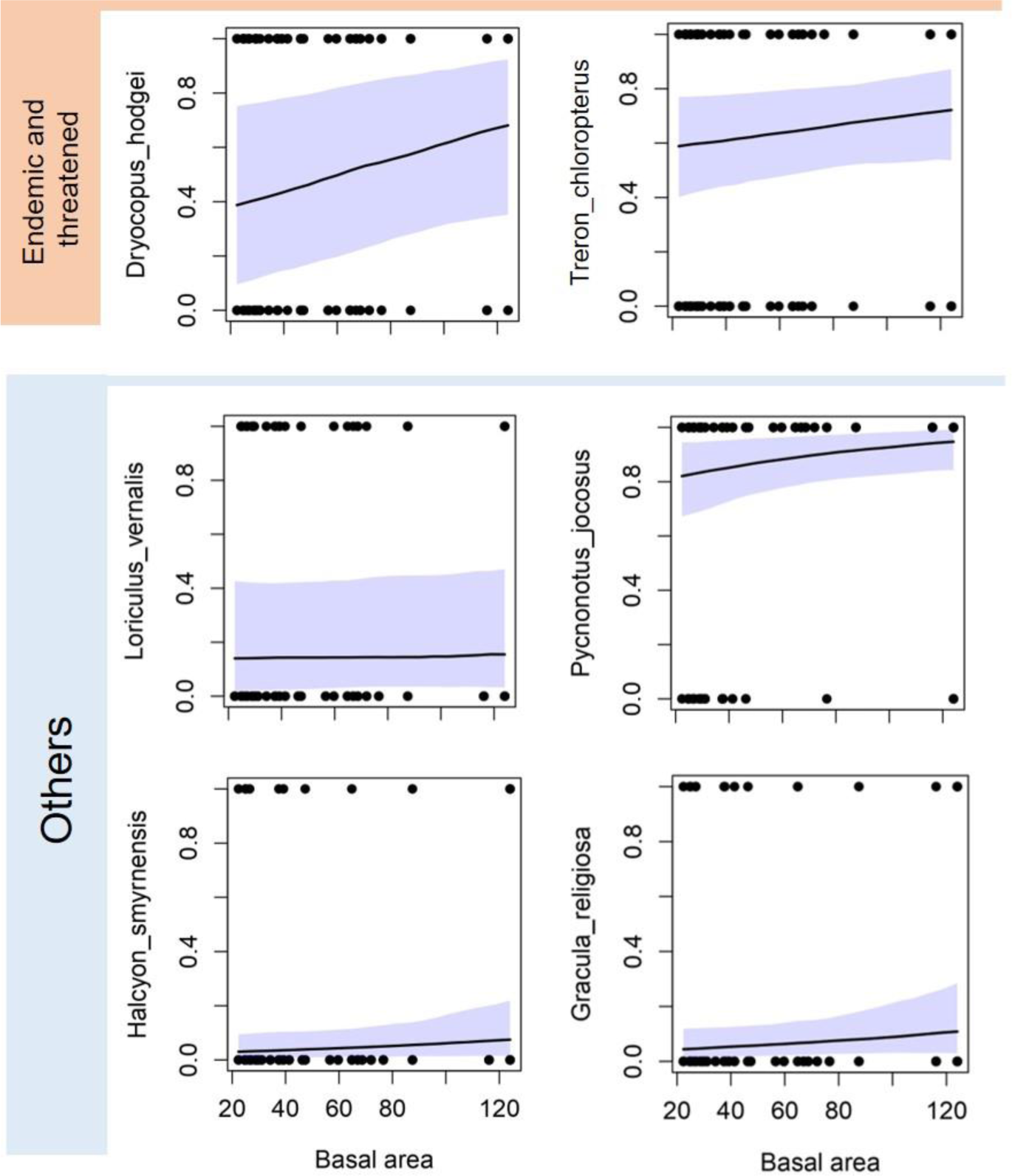
HMSC model predictions showing changes in bird species occurrence probabilities (y-axis) with basal area. Only statistically detectable relationships have been shown. Black lines represent the posterior median, points represent individual samples, and the shaded area depicts the posterior 95% credible interval. Species names are provided along the y-axis of each plot.

**Figure S6.**
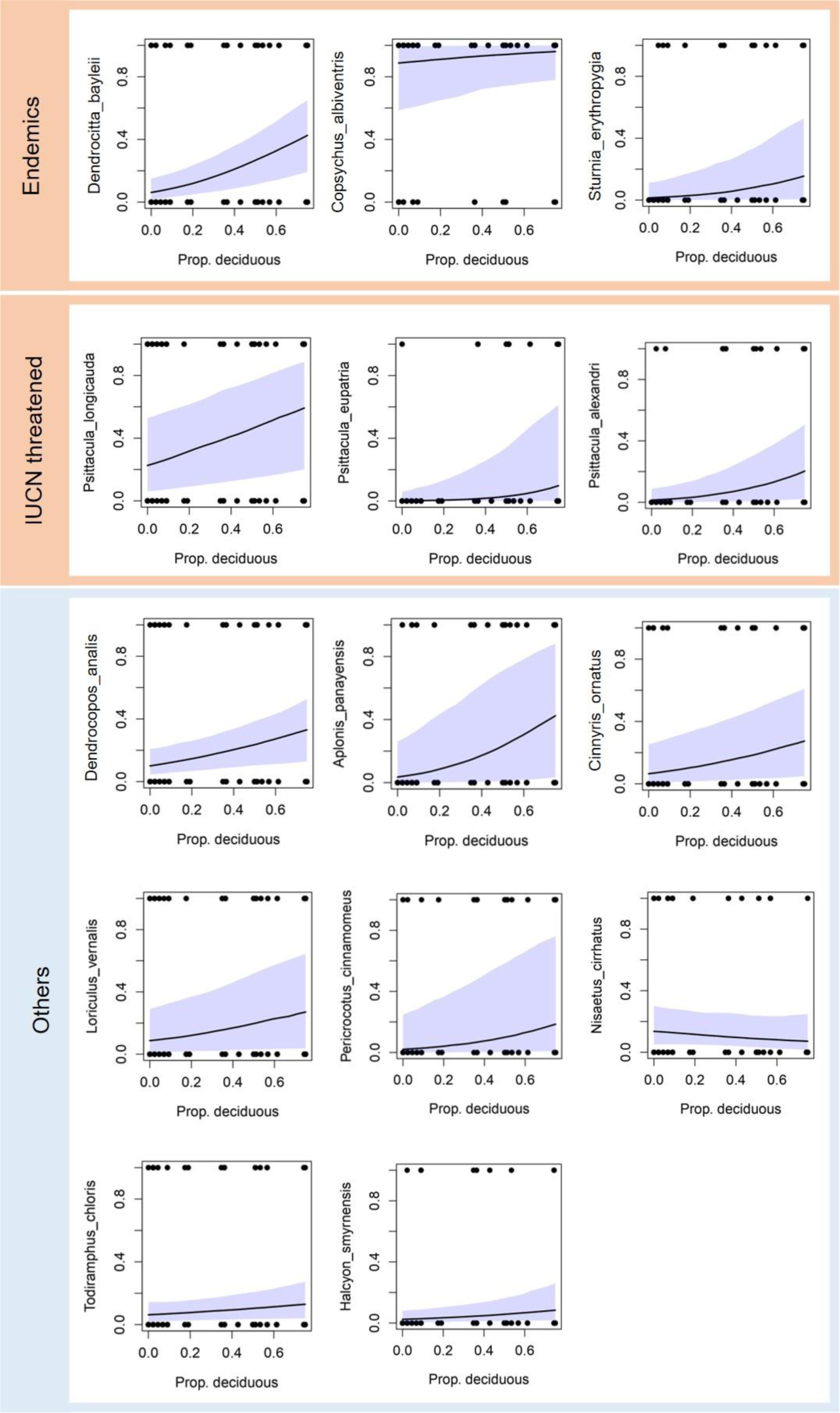
HMSC model predictions showing changes in bird species occurrence probabilities (y-axis) with proportion of deciduous trees. Only statistically detectable relationships have been shown. Black lines represent the posterior median, points represent individual samples, and the shaded area depicts the posterior 95% credible interval. Species names are provided along the y-axis of each plot.

**Figure S7.**
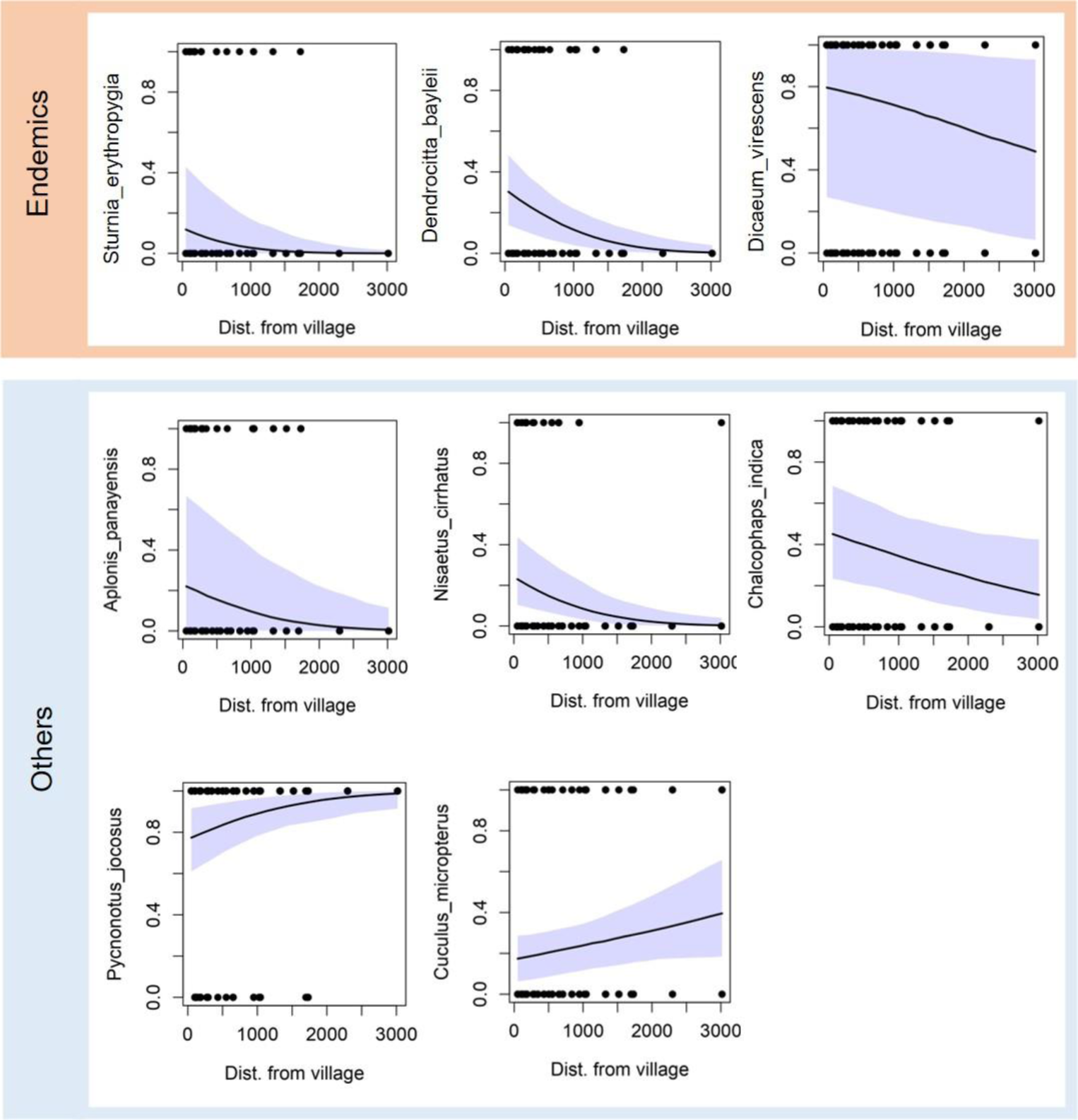
HMSC model predictions showing changes in bird species occurrence probabilities (y-axis) with increasing distance from nearest settlement/village. Only statistically detectable relationships have been shown. Black lines represent the posterior median, points represent individual samples, and the shaded area depicts the posterior 95% credible interval. Species names are provided along the y-axis of each plot.

